# An Aerobic Anoxygenic Phototrophic Bacterium Fixes CO_2_ via the Calvin-Benson-Bassham Cycle

**DOI:** 10.1101/2021.04.29.441244

**Authors:** Kai Tang, Yang Liu, Yonghui Zeng, Fuying Feng, Ke Jin, Bo Yuan

## Abstract

Aerobic anoxygenic phototrophic bacteria (AAnPB) are photoheterotrophs, which use light as auxiliary energy and require organic carbon (OC) for growth. Herein, we report the unusual strain B3, which is a true AAnPB because it requires oxygen for growth, harbours genes for *cbb3*- and *bd*-type cytochromes and *acs*F, and produces bacteriochlorophyll. The B3 genome encodes the complete metabolic pathways for AAnPB light utilization, CO_2_ fixation via Calvin-Benson-Bassham (CBB) cycle and oxidation of sulfite and H_2_, and the transcriptome indicated that all components of these pathways were fully transcribed. Expression of the marker genes related to photosynthesis, including *puf*M for light harnessing and *rbc*L for CO_2_ fixation, and the activity of RubisCO, the key enzyme in the Calvin-Benson-Bassham (CBB) cycle, increased in response to decreased OC supply. Large amounts of cell biomass were obtained in liquid BG11 medium under illumination. The strain thus likely photoautotrophically grows using sulfite or H_2_ as an electron donor. Similar GC contents between photosynthesis, the CBB cycle and 16S rRNA genes and the consistency of their phylogenetic topologies implied that light harnessing and carbon fixation genes evolved vertically from an anaerobic phototrophic ancestor of Rhodospirillaceae in Alphaproteobacteria. In conclusion, strain B3 represents a novel AAnPB characterized by photoautotrophy using the CBB cycle. This kind of AAnPB may be ecologically significant in the global carbon cycle.

## Introduction

Aerobic anoxygenic phototrophic bacteria (AAnPB) are obligate aerobes and perform photosynthesis without producing oxygen [1, 2]. They use the pigment bacteriochlorophyll (BChl) and carotenoids to harvest solar energy and the type II reaction centre (RCII) to photochemically perform electron transfer and eventually drive ATP biosynthesis [1, 3]. Light may promote the growth of AAnPB. For instance, the utilization of light energy prolonged the survival of *Dinoroseobacter shibae*, a major marine AAnPB, under organic carbon (OC) starvation [4, 5]. A freshwater AAnPB strain, *Aquincola tertiaricarbonis* L108, was induced to produce BChl *a* and used light as an auxiliary energy source to maintain its growth only when fed at an extremely low substrate rate [6]. In addition to its reported promotion of the growth of pure cultured isolates, light was also found to enhance the growth rate of AAnPB species in natural environments [7, 8]. The growth promotion by supplementary light resulted from the absorption of light energy reducing the respiration consumption of OC for energy production, which increased the nutrient supply for the anabolism of AAnPB when OC was less available [9-11]. Therefore, phototrophy makes AAnPB more competitive and dominant than strict heterotrophs in the open ocean, where there is little organic matter available [12].

Carbon fixation provides the basis for all life on Earth [13], and seven carbon fixation pathways have evolved in nature [14]. Among the seven pathways, the Calvin-Benson-Bassham (CBB) cycle is the most widespread [15] and responsible for the most carbon fixation on Earth [16]. The key catalyst of this fixation is the enzyme RubisCO, which is composed of a large subunit encoded by the *rbc*L gene and in some cases a small subunit encoded by the *rbc*S gene. Photoautotrophic bacteria, including aerobic cyanobacteria and some obligate or facultative anaerobic anoxygenic phototrophs, use the CBB cycle for CO_2_ fixation, enabling autotrophic growth [17, 18]. The first clue to AAnPB harbouring a complete CCB pathway was a recent marine metagenomics analysis that revealed that the assembled genome of a globally distributed AAnPB possessed both *puf*M and *rbc*L/S genes, demonstrating great potential for primary productivity [19]. However, whether the genes associated with solar energy utilization and carbon fixing are expressed and how the related physiological activities are performed still need to be ascertained through multiple assays, especially those based on pure cultured bacterial strains. Additionally, evidence was found that some AAnPB showed the physiological activities for inorganic carbon fixation in other carbon fixation pathways, but that was not substantial enough to support autotrophic growth. For example, *Roseobacter* clade bacteria could utilize carbon monoxide as an additional energy or carbon source [11, 20], and *Dinoroseobacter shibae* fixed CO_2_ using the ethylmalonyl-CoA (EMC) pathway [21].

In arid and semiarid areas, biological soil crusts (BSCs) are widely distributed as one of the major features of the Earth’s terrestrial surface and play a vital role in the global carbon cycle [22]. Benefiting from supplementary light, AAnPB may occupy an important niche in such organic substrate-poor terrestrial environments [23, 24]. Moreover, our previous work showed that AAnPB could greatly enhance organic matter content and promote the development of BSCs [25]. For these reasons, the contribution of carbon accumulation by AAnPB was hypothesized to be significant in BSCs.

In this study, we used the spread and streak plate technique to isolate AAnPB strains from BSCs. Genome and transcriptome analyses revealed that one AAnPB strain, B3, possessed the genes for a full CCB cycle pathway and that all the components in this pathway were transcribed. Physiological assays further ascertained that the strain produced the enzyme RubisCO with high physiological activity and was photoautotrophic with high biomass in media with little or no OC added. This will shed light on the evolution of AAnPB and their role in the global carbon cycle.

## Materials and Methods

### Sampling and isolation

Moss-dominated biological soil crusts (BSCs) (about 2 cm of top soil) were collected in October 2016 from sand dunes (42.427 N, 116.769 E, 1380 m asl) located in the southeastern Hunshadake Desert in China. The sampled soils were stored in aseptic sampling bags in a vehicle-mounted refrigerator and brought to the laboratory within 72 hours for further treatment.

Bacteria were isolated on 1/2-strength R2A agar (Table S1) using standard tenfold dilution plating and streak cultivation techniques. The plates were incubated aerobically at 28 °C under a light regimen of 4000 lux with a 12/12 hour cycle (light/dark).

### Identification of AAnPB strains

Bacterial strains possessing the *puf*M gene, encoding the M subunit of the type II reaction centre, were tentatively considered AAnPB. The presence of the *puf*M gene was confirmed through PCR and sequencing. The PCR template for *puf*M amplification was prepared using a rapid cell lysis protocol for which colonies harvested from plates were directly incubated in 0.05 M NaOH with 95/4 °C cycling. The 27F/1492R and *puf*M_557F/*puf*M_WAW primer pairs (Table S2) were used to amplify 16S rRNA and *puf*M genes, respectively, and PCR products were sequenced using the Sanger sequencing method by Sangon Biotech Comp, Shanghai. The absorption spectra of *puf*M-positive isolates *in vivo* were recorded on a Synergy H4 Hybrid Reader (BioTek) with a scanning range from 300 to 900 nm. Bacteriochlorophyll pigments were extracted from cells grown in 1/10-strength R2A liquid medium with pre-cooled acetone/methanol (7:2) for 2 hours and determined using HPLC as described previously [25]. BChl *a* standard was purchased from Sigma-Aldrich Fine Chemicals.

### Tests of oxygen requirement for growth

Strain B3 was incubated on 1/2-strength R2A agar in an anaerobic atmosphere by using MGC AnaeroPack-JAR and MGC AnaeroPack-Anaero (Mitsubishi Gas Chemical Company, Tokyo, Japan) (Figure S1a) and agar shake tube technique [26] on Medium A [27] (Figure S1b). The culture was conducted at 28 °C under illumination (4000 lux of luminous intensity, 12 h/12 h light-dark cycle) and no illumination (in dark) (Figure S1b). One facultative anaerobic strain, *Rhodocista centenaria* DSM9894^T^ [28] was used as a reference, and three replicates were set up in these tests of oxygen requirement for growth.

### Photoautotrophic growth tests

Cell biomass (fresh and dry cell weights) was used to determine the photoautotrophy of AAnPB strains grown in BG11 liquid media (Table S1). Cells grown in 1/10 R2A medium were centrifuged, washed three times and resuspended in NaCl solution (0.08%). The cell suspension (approximately 1.00 of OD_600 nm_) of 100 µL was inoculated into 50 mL of BG11 liquid medium with variant trace levels of vitamin B12 to incubate in a shaker at 180 r/min under illumination (4000 lux of luminous intensity, 12 h/12 h light-dark cycle). After five days of incubation, the cells were collected by centrifugation. The centrifuge tubes were placed upside-down and air-dried for 2 hours, and the fresh cell biomass was determined. Dry cell weights were obtained after drying at 80 °C for 24 hours. The levels of vitamin B12 were 0, 0.2, 0.4 and 1.0 μg per 50 mL of medium. Five repeats were carried out.

Total bacterial numbers of strain B3 were used to determine the growth curve of AAnPB strains in BG11 liquid media under illumination and dark. The pH was adjusted to 7.2, 1.0 μg per 50 mL vitamin B12 was added, and the sample was then incubated in a shaker at 180 r/min under illumination (4000 lux of luminous intensity, 12 h/12 h light-dark cycle). Cell numbers were counted by hemacytometer (Qiujing XB-K-25, Shanghai, China) under a phase contrast microscope (Nikon ECLIPSE Ti, Japan), and the following formula was used:

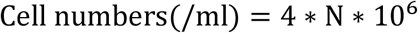

the letter “N” refers to the cell number in one chamber unit.

### Treatments of different organic carbon levels

Based on R2A, four OC treatments (labelled 5A, 2.5A, A and 0A, respectively) (Table S1) were set up to determine the effect of OC on photosynthetic gene expression and RubisCO enzyme activity. Cells were cultivated in a thermostatic oscillator at 180 r/min at 28 °C under 4000 lux with a 12/12 hour (light/dark) cycle; then cells at the early steady growth phase were harvested by centrifugation at 12,000 × g under 4 °C for 15 min.

### RubisCO enzyme activity assay

The RubisCO enzyme activities at different OC levels were measured in a reaction mixture (200 μl) containing the buffer, substrates, cofactor and enzymes prepared according to Takai et al. [29] and measured by spectrophotometry as described by Yuan *et al*. [30]. Briefly, 200 μl of reaction liquid was transferred to a 96 micro-well plate, and the absorption at 340 nm was measured using the Synergy H4 Hybrid Reader (BioTek). Then, after 0.1 mL of 25 mM ribulose-1,5-bisphosphate (RuBP) was added to the well and reacted for 30 s, the absorption at 340 nm was measured again. For the control treatment, no RuBP was added. To calculate RubisCO’s enzyme activity, the following formula was used:

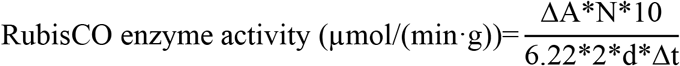

where ΔA is the change in OD_340 nm_ during 30 s the change in the control, the letter “N” refers to the dilution ratio (100-fold used in this study), 6.22 is the absorbency coefficient of 1 mm NADH at OD_340 nm_, Δt is the reaction time (30 s in this study) and the letter “d” represents the colorimetric optical path, which was 0.0635 cm in this study.

### ^13^CO_2_ assimilation assay

First, 1.59 mg/L NaH^13^CO3 (98 atom% ^13^C, Sigma-Aldrich, USA, no. 372382) was added to liquid BG11 medium to test the CO_2_ assimilation ability of strain B3. Then 5.0 ml of strain B3 BG11 medium (OD=0.03) was added into 4.0 L of liquid BG11 medium containing NaH^13^CO3, which was then cultured with magnetic stirrers (IT-07A3, Shanghai Yi Heng Co., Ltd, Shanghai, China) at 180 r/min at 28 °C under a light regimen of 4000 lux and a 12/12 hour (light/dark) cycle. After 120 hours of culture, the cells were acquired by centrifuged at 12,000 × g for 10 min, washed 3 times with 10^−3^ M H_3_PO_4_ (pH=3.0), and lyophilized using a freeze dryer (Christ Alpha 1-2 LD Plus, France).

The ^13^C contents in cell matter (after combustion to CO_2_) and in the CO_2_ gas used as a carbon source were determined by a Thermo Scientific MAT 253 plus mass spectrometer fit with a Gasbench II-IRMS at Beijing Zhong Ke Bai Ce Testing Technology, China. The δ^13^C value was presented as per mille deviation from the Vienna Pee Dee Belemnite standard (VPDB) [31, 32] as follows:

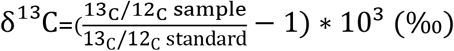

### Whole-genome sequencing and analysis

The genomic DNA of the B3 strain was extracted using a Bacterial Genomic DNA Purification Kit (DP302-2, Tiangen Biotech Co., Beijing, China) according to the manufacturer’s protocols. Genome sequencing was performed on the PacBio RSII platform at Shanghai Majorbio Pharm Technology Co., Ltd. After filtering the low-quality reads and adapters, clean reads were assembled into a single gap-free contig using HGAP ver 3.0 [33]. The open reading frames (ORFs) of the assembled genome were predicted with Glimmer ver. 3.02 [34]. The genome was annotated by the Kyoto Encyclopedia of Genes and Genomes (KEGG) website (http://www.genome.jp/kegg/) [35] and visualized with Circos software [36]. SWISS-MODEL was used to model the RubisCO enzyme structure online based on *cbb*L/S/X genes online (https://swissmodel.expasy.org/interactive) [37].

### Assays for gene expression quantification and transcriptomics

Cells were grown in media with different levels of OC sources (Table S1), and the expression of genes for light utilization and carbon fixation was determined and compared. Cells were cultivated in a thermostatic oscillator at 180 r/min and 28 °C under natural light for five days and then collected through centrifugation at 12,000 × g for 15 min at 4 °C.

Total cellular RNA was extracted using the RNAprep Pure Cell/Bacteria Kit (DP430, Tiangen Biotech Co, Beijing). RNA integrity was examined through agarose gel electrophoresis, and RNA purity was determined with a NanoDrop spectrophotometer (Thermo Fisher Scientific). After pretreatment with RNase-free DNase I (TaKaRa, Dalian, China), total RNA (1 μg) was used to synthesize first-strand cDNA by employing an M-MLV Reverse Transcriptase (code no. 2641A, TaKaRa Bio Inc., Dalian, China) according to the manufacturer’s instructions. The cDNA was employed as a template using SYBR Premix Ex Taq II (Takara) in a LightCycler 480 Real Time PCR system (Roche, Basel, Switzerland) for qRT-PCR analysis, including the 16S rRNA, *puf*M and *rbc*L genes. The primers and thermal programs are described in Table S2. The standard curve of the 16S rRNA gene was created as previously described [25]. The 16S rRNA gene was employed as a reference gene to normalize the expression levels of target genes in samples, and the 2^-ΔΔCT^ method [38] was used to calculate the relative expression levels of the target genes. Three technical replicates of qPCR were performed for each reaction.

Total RNA sequencing was conducted by Shanghai Majorbio Pharm Technology Co., Ltd. (Shanghai, China) on a HiSeq Illumina platform. RNA-seq reads were trimmed using SeqPrep (https://github.com/jstjohn/SeqPrep) and Sickle (ver. 1.33). Diamond (ver. 0.8.35) (https://github.com/bbuchfink/diamond) was used to ensure that the rRNA ratio was lower than 5%. Based on the Burrows-Wheeler transform, Bowtie 2 (ver. 2.2.9) was used to align trimmed clean reads [39]. The relative expression in transcripts per million reads (TMP) was estimated by Salmon (ver. 0.8.2) (https://github.com/COMBINE-lab/salmon). ORFs were annotated with the KEGG database.

### Assays for putative electron donors for carbon fixation

After the surveys of the genome and transcriptome, potential electron donors were listed and then added to cultures to observe their effects on the biomass. Compounds markedly enhancing cell biomass were probable electron donors [40]. Na_2_SO_3_ and H_2_ were selected and further assayed, respectively. Na_2_SO_3_ (48 mg/L) was added into liquid BG11. H_2_ was filled as headspace gas (20% H_2_ + 80% air). The cultivation conditions, cells harvesting and biomass (cells dry weight) were performed as described above.

### Phylogenetic analyses

Phylogenies of the 16S rRNA, photosynthetic and carbon fixation genes were constructed and compared to reveal the evolution of AAnPB and their photosynthetic apparatus. The phylogenetic tree of phototrophic bacteria [41] was referred to to obtain reference sequences of the 16S rRNA gene from EzBioCloud (www.ezbiocloud.net/). The *puf*LMC-*bch*XYZ reference amino acid sequences were recovered from Imhoff *et al*. [41]. Reference amino acid sequences of *rbc*L were obtained from the NCBI (nonredundant protein database, *nr*) through BLASTP ver. 2.10.0+ [42]. Multiple alignments were performed with the MUSCLE program, and the maximum-likelihood (ML) phylogenetic trees were calculated using MEGA ver. X with 1,000 tree resamplings. The Kimura 2-parameter model (16S rRNA gene) [43] and Jones-Taylor-Thornton (JTT) amino acid substitution model (*puf*LMC-bchXYZ and *rbc*L gene) [44] were used. The phylogenetic trees were visualized using iTOL software online.

### Statistical analyses

Significant differences were calculated using Tukey’s test and identified at P < 0.05 by GraphPad Prism 8.0.

## Results and Discussion

### Isolate B3 is a true AAnPB of Alphaproteobacteria

A pink-pigmented bacterial isolate, designated B3, was obtained on 1/2 strength R2A medium from moss-dominated BSCs, using aerobic spread and streak plates. No growth of B3 was observed on 1/2 R2A agar plates under anaerobic conditions, even for a longer culture time (31 days) (Figure S1a). Under either illumination or no illumination, the strain’s growth was obvious at only the top of the tube (Figure S1b). This was distinct from the growth of the reference strain *Rhodocista centenaria* DSM9894^T^, which occurred in the whole tube with Medium A. *Rhodocista centenaria* DSM9894^T^ was identified as a facultative anaerobic purple non-sulphur bacterium (PNSB) [28]. In Medium A, PNSB were characterized by growing in anaerobic conditions under illumination [27]. These growth tests showed that strain B3 is obligate aerobic. This was also supported by the presence of microaerobic *ccb*_*3*_-type and highly-aerobic *bd*-type cytochrome oxidases and oxygen-dependent ring cyclase (*acs*F) revealed through the strain’s genome survey and gene PCR and sequencing (Table S3). Anaerobic phototrophs distinctly lack aerobic cytochrome oxidases especially *ccb*_3_-type [45] and *bd*-type [46] cytochrome oxidases. Gene *acs*F encodes the aerobic form of Mg protoporphyrin IX monomethyl ester oxidative cyclase necessary for bacteriochlorophyll (BChl) biosynthesis in oxic environments [47, 48], and therefore, the genome’s possession if *ccb*_3_- and *bd*-type cytochrome oxidase and *acs*F genes was used as key evidence for ‘*Candidatus* Luxescamonaceae’ having an aerobic nature and thus being a true AAnPB [19].

The viable cells of strain B3 grown with vitamin B12 (VB12) supplied showed an absorption peak at 870 nm (Figure 1a), which is a typical feature of type II reaction centres in AAnPB [6, 12]. The HPLC elution profile of the pigment extracts showed a peak at a retention time of 21 min with peak absorption at 710 nm identical to that of standard BChl *a* (Figure 1b). This validated that strain B3 could produce BChl *a* under aerobic conditions. However, without supplementation with VB12, BChl *a* was not detected. VB12 regulated photosystem gene expression to affect the production of Bchl *a* via the *Crt*J antirepressor *Aer*R [49]. In BSCs, cyanobacteria and algae are commonly dominant microbes [50] and common VB12 producers [51]. When we cocultured strain B3 with *Microcoleus vaginatus*, a cyanobacterium dominant in BSCs, Bchl *a* production was detected (Figure S2). PCR amplification and sequencing results demonstrated that strain B3 possessed the *puf*M gene, encoding the M subunit of the type II photosynthetic reaction centre in AAnPB and, the sequence was closely related to that of *Rhodocista centenaria* (similarity of 83.75%).

**Figure 1.**
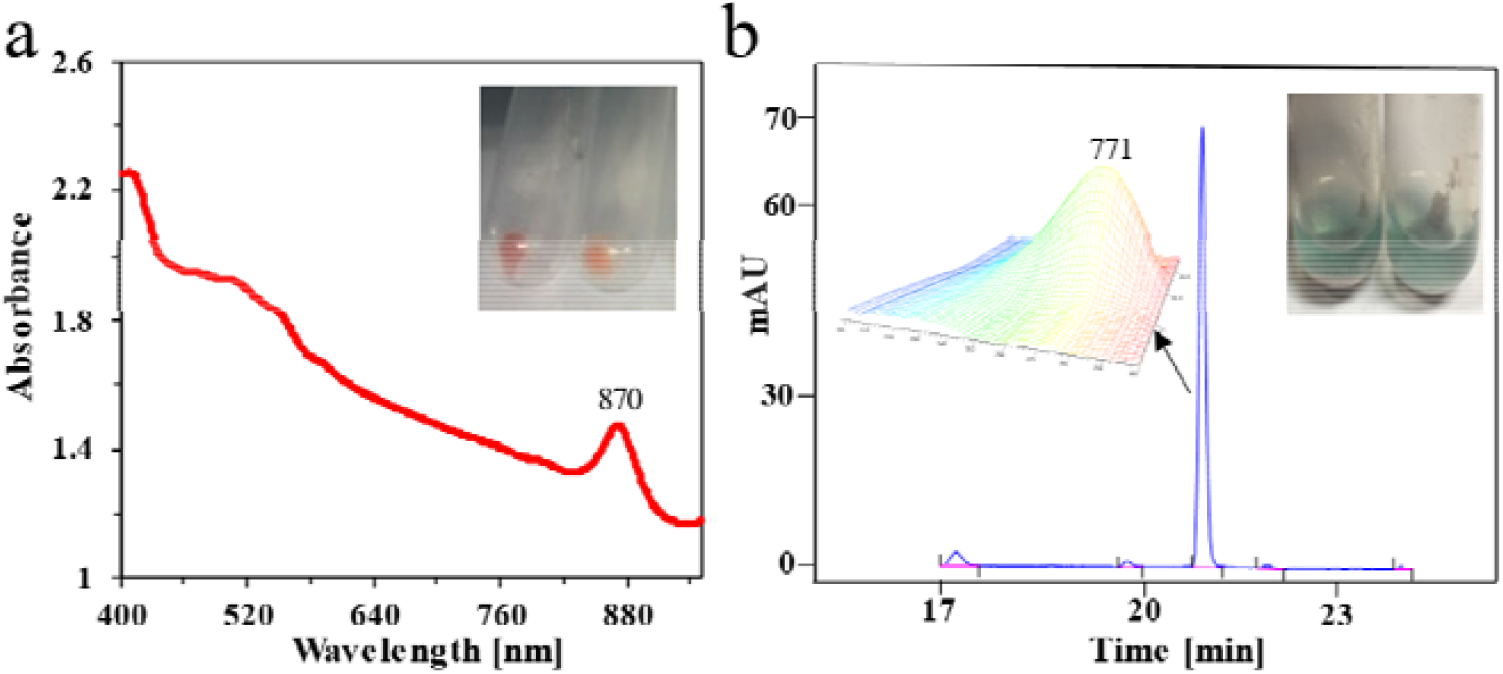
Cell and pigment characterization of strain B3. (a) Absorption spectrum of viable cells between 300 - 900 nm; the top right shows the cell pellets. (b) Chromatogram of extracted pigments; the top right shows the extract of the pigments.

The 16S rRNA gene of strain B3 shared its highest similarity of 93.09% with that of *Oleisolibacter albus* NAU-10^T^, which was a strictly aerobic species belonging to of the family Rhodospirillaceae of the class Alphaproteobacteria [52], suggesting that strain B3 belongs in Alphaproteobacteria.

Collectively, these results strongly suggest that strain B3 is a true AAnPB of Alphaproteobacteria.

### Strain B3 grew photoautotrophically

AAnPB can harvest light to obtain additional energy to supplement their heterotrophic growth [6, 53]; some AAnPB strains can even fix CO or CO_2_, but the fixed carbon cannot support autotrophic growth without a substantial OC supply [12, 20, 21]. Under illumination, strain B3 grew in liquid BG11 media in the presence of trace citrate (6 mg/L) and VB12 (0, 0.2, 0.4 and 1.0 μg per 50 mL) without other OC addition (Figure 2a). At 0.4 μg/50 mL VB12, the biomass of fresh and dry cells peaked at 110 mg and 8 mg per 50 mL, respectively. The production of newly synthesized biomass from CO_2_ is a sign of carbon autotrophic microbes [54]. Moreover, the cells grown with ^13^CO_2_ exhibited δ^13^C values of 24.71 - 30.65 %, and this directly supported strain B3 having the ability to fix CO_2_ under illumination (Table 1). Cultivated in BG11 media under illumination, strain B3 showed a typical growth curve of single-celled bacteria with a lag phase of 5 hours and logarithmic phase of 50 hours; in contrast, under dark, the strain cell numbers were only slightly increased after 80 hours (Figure 2b). These suggested that strain B3 could grow photoautotrophically. Photoautotrophy may make strain B3 more competitive in OC-deficient BSCs. In fact, the inoculation of strain B3 effectively promoted the development of BSCs and the soil OC heavily accumulated [55], suggesting that AAnPB may make great contributions for carbon sequestration in arid land. High biomass yield through carbon fixation of CO_2_ may help solve the global warming problem [56]. Metagenomic analysis indicated that AAnPB was a potential significant contributor to the carbon cycle in marine environment, because they may widely distributed in CBB at high abundance [19]. Therefore, photoautotrophic AAnPB may be ecologically important worldwide.

**Table 1.**
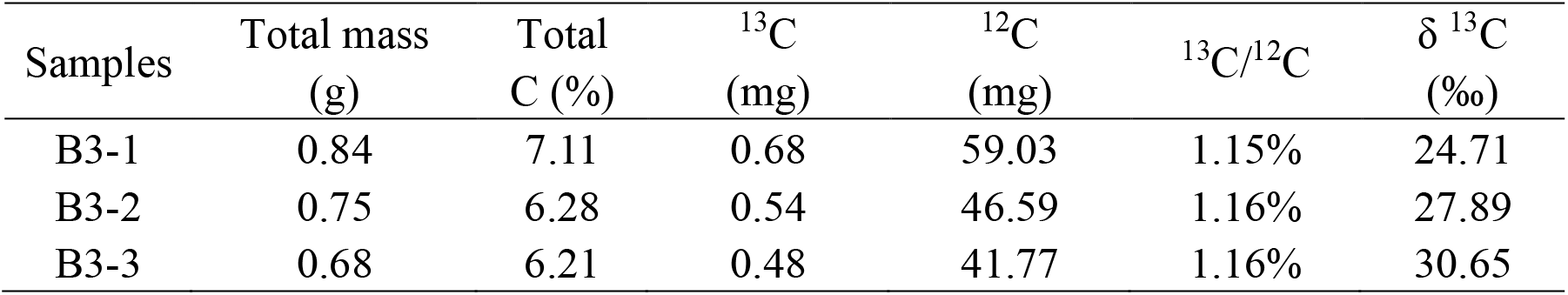
δ^13^C ratios in dried strain B3 cells.

**Figure 2.**
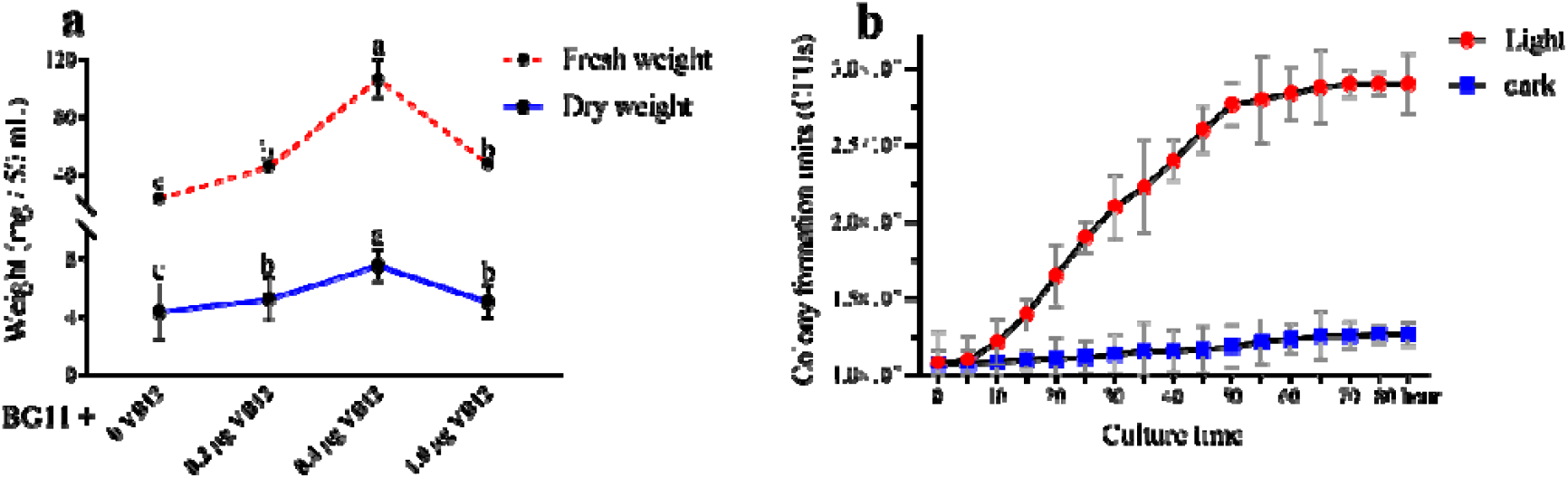
Biomass of strain B3 in 50 mL of liquid BG11 with different levels of added VB12 (a), and the growth curve of strain B3 in liquid BG11 under illumination and dark (b). Letters (a, b, or c) in the superscript represent the observed significant differences (Tukey’s test; P < 0.05).

### Genome of strain B3 contains complete pathways of the type II reaction centre for light utilization and the CBB cycle for CO_2_ fixation

The whole genome of strain B3 was sequenced to explore the metabolic pathways responsible for its photoautotrophic growth, and the GenBank accession number is CP051775. The results showed that the genome of strain B3 contained one chromosome with a length of 3664154 bp and a GC content of 69.26 % and four plasmids (Figure 3a; Table S4).

**Figure 3.**
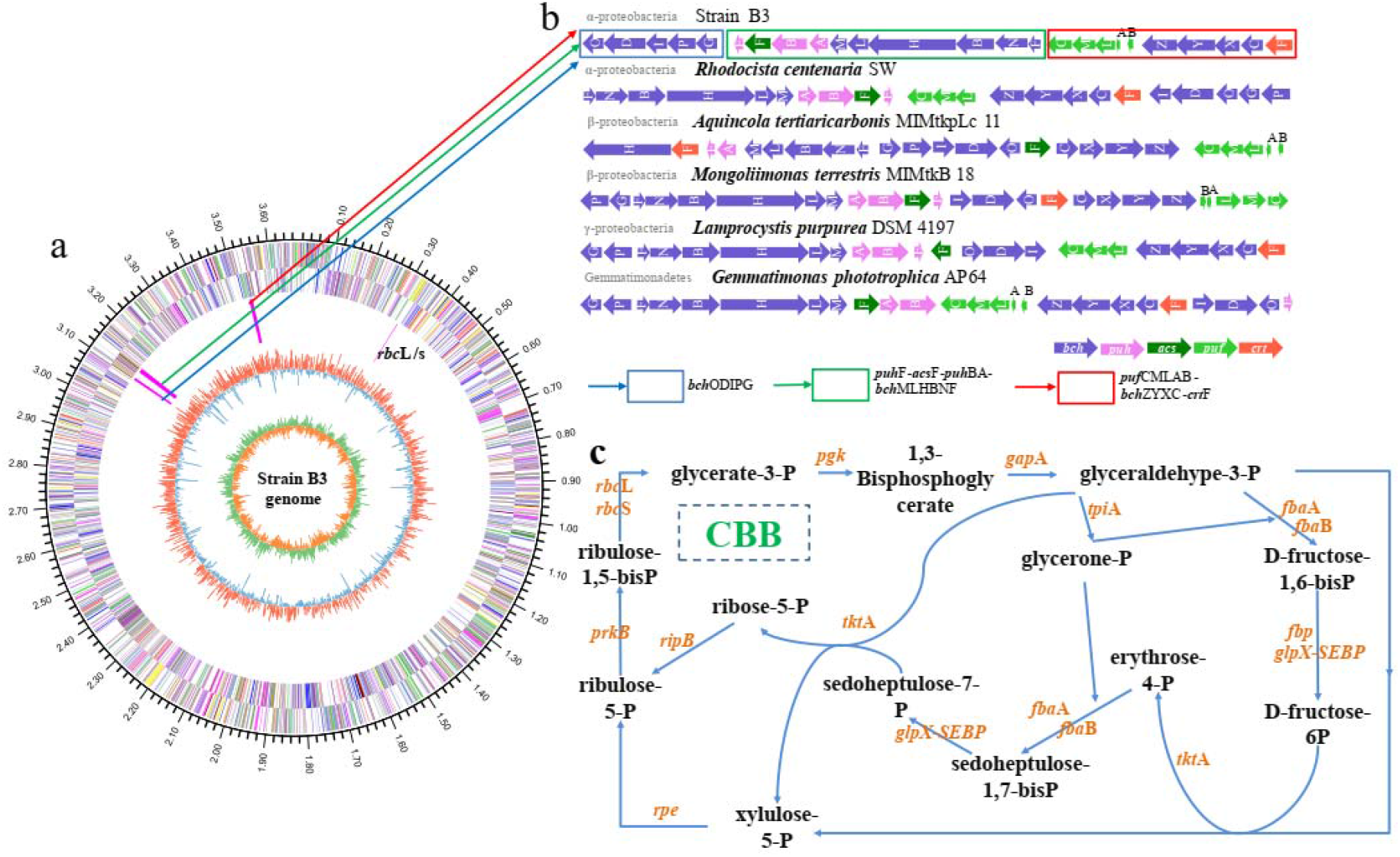
Genome, photosynthesis gene clusters and carbon fixation metabolism of strain B3. (a) Circular genome of strain B3. The first outer circle shows the size of the genome; the second and third are the coding sequences (CDSs) with different colours denoting various functions based on COG; the fourth circle contains photosynthesis gene clusters (PGCs) and *rbc*L/S. PGCs are distributed in three positions in the genome. The fifth circle is the GC content, and the red and blue peaks represent higher or lower GC content than average; the inside is the value of GC-skew, and green and orange represent the leading and lagging strands, respectively. (b) The PGCs in 6 AAnPB from Alpha-, Beta-, and Gammaproteobacteria and Gemmatimonadetes. (c) Overview of the metabolic pathway of the Benson-Bassham-Calvin cycle (CBB).

The photosynthetic genes clustered into a photosynthesis gene cluster (PGC), which is a typical feature in most purple phototrophic bacteria [57], heliobacteria [58] and Gemmatimonadetes [48]. The PGC identified in the B3 genome is 44.3 kb long and consists of genes in the order *bch*ODIPG, *puh*F-*acs*F-*puh*BA-*bch*MLHBA and *puf*CMLAB-*bch*ZYXC-*crt*F (Figure 3b), similar to those of proteobacterial AAnPB [59]. As the presence of *puf* genes, which encode bacterial reaction centre subunits, and genes necessary for bacteriochlorophyll biosynthesis in the PGC suggests, strain B3 may carry out a phototrophic strategy with type II reaction centres.

To date, there are seven known pathways for carbon fixation, including the CBB cycle, reductive tricarboxylic acid (rTCA) cycle, 3-hydroxypropionate bi-cycle (HBC), Wood-Ljungdahl (WL) cycle, dicarboxylate/4-hydroxybutyrate (DH) cycle, 4-hydroxypropionate cycle, and reductive glycine pathway [14]. None of pathways except CBB has been identified in strain B3. Strain B3 assimilates CO_2_ through the CBB cycle, which is complete in the genome (Figure 3c). RubisCO is a key enzyme of the CBB cycle, in which CO_2_ serves as a substrate in organisms evolved from a non-CO_2_-fixing and non-autotrophic ancestor [60]. The strain B3 genome possesses both an *rbc*L and an *rbc*S gene, encoding large and small subunits of RubisCO, respectively. The *cbb*X gene encoding RubisCO activase, similar to that of *Rhodobacter sphaeroides* [61], was also identified in B3. Additionally, a complete pathway of crassulacean acid metabolism (CAM) is present in B3 (Table S3), which is another possible mechanism for the concentration of CO_2_, in addition to that through the small subunits of RubisCO, and it is widely employed in C4 plants and some algae [62].

### Pathways of light utilization and CO_2_ fixation are active in strain B3

To confirm whether the photosynthetic pathways for light utilization and CO_2_ fixation are active, we determined the shifts in the transcriptome, the physiological activity of the enzyme RubisCO and the quantitative expression levels of the *puf*M and *rbc*L genes in response to varying OC concentrations. Transcriptome analysis showed that genes involved in the bacteriochlorophyll-dependent light utilization of the type II reaction centre, CO_2_ fixation of the CBB cycle and CO_2_ condensation of CAM were all transcribed at different levels (Figure 4). The expression levels of both *puf*M and *rbc*L genes and RubisCO enzyme activity were significantly enhanced in strain B3 when the OC concentration greatly decreased (Figure 5). The increased expression in response to the OC limit implied that the AAnPB strain B3 modified its energy and carbon source strategy from heavily relying on chemical energy from OC to relying more on optical energy and CO_2_ fixation via the CBB cycle. The strategy of light utilization was commonly adopted by other AAnPB as well [6- 8, 10, 63, 64], and OC starvation drives a few AAnPB strains to assimilate CO_2_ [21] or CO [29, 65], but carbon fixation via the CBB cycle has not been reported. Moreover, the *puf*M and *rbcL* gene copy numbers were greatly correlated with cell density (r=0.99, P=0.0034; r=0.80, P=0.198) and cell biomass (r=0.97, P=0.0318; r=0.89, P=0.108) (Table S5). The strong correlation of photosynthetic genes and growth features suggested that light utilization and CO_2_ fixation were connected in this strain. This may be explained as light energy is primarily consumed by CO_2_ fixation through the CBB cycle in microbial photoautotrophy [66, 67].

**Figure 4.**
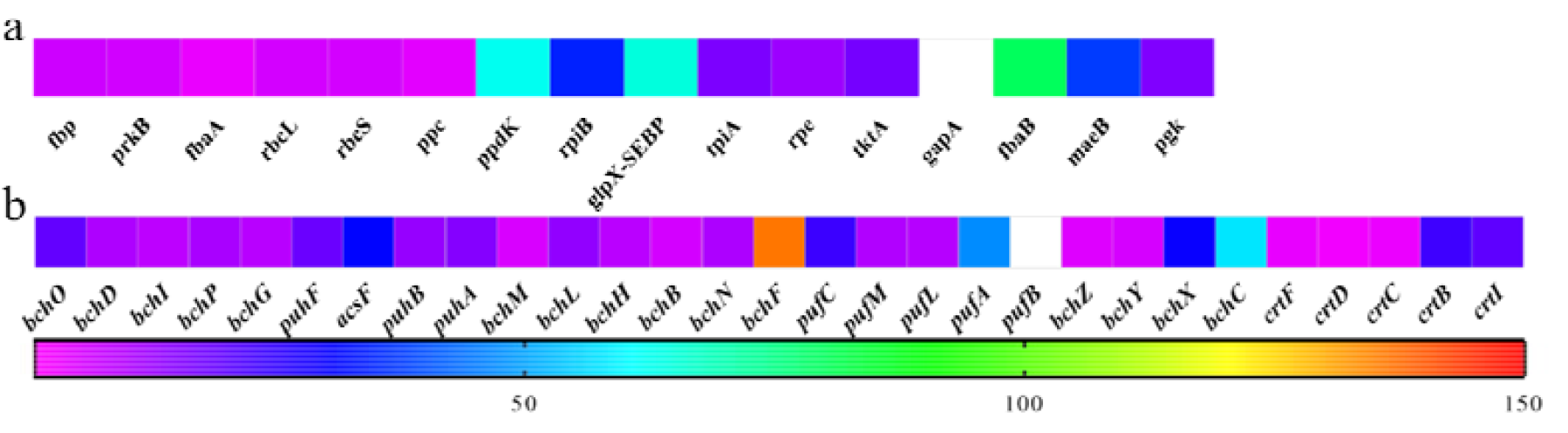
Heatmap of the expression of genes involved in carbon fixation (a) and photosynthesis gene clusters (PGCs) after five days of culture at an OC concentration of 2.5A (b), including the *bch, puf, puh*, and *crt* operons and the *acs*F gene. The values of the *gap*A and *puf*B genes are outside the defined range (TPM values are 201.54 and 386.06, respectively) and are represented in white. Transcripts per million reads (TPM value) were used to represent differences in gene expression.

**Figure 5.**
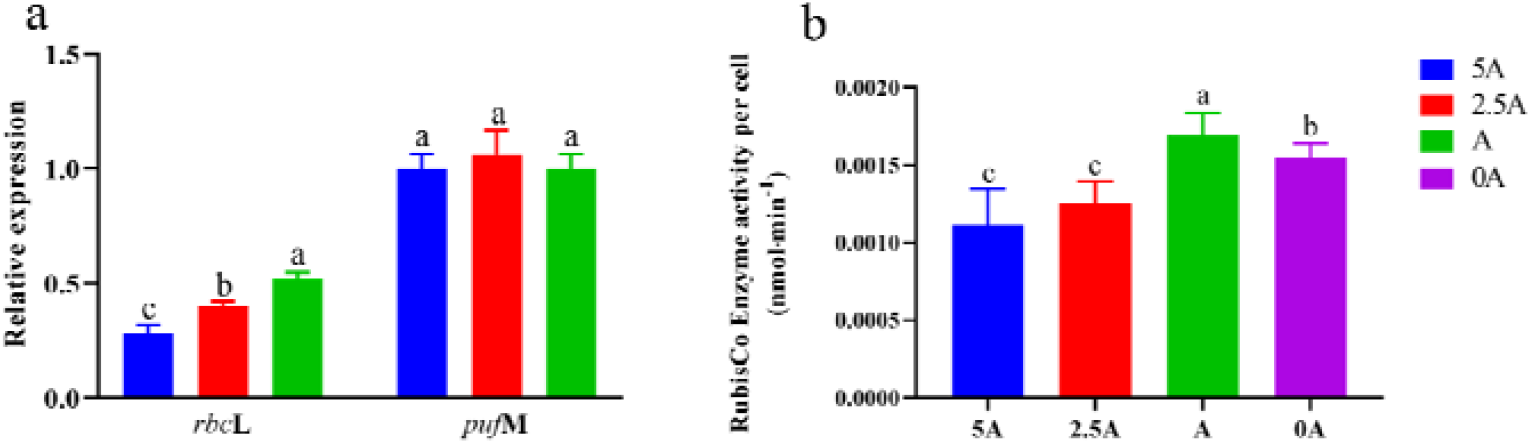
Relative gene expression and RubisCO enzyme activity changes in strain B3 at different organic carbon levels. The relative expression of *puf*M and *rbc*L genes (a), RubisCO enzyme activity per cell; (b) changes in different organic carbon levels, including 5A (blue), 2.5A (red), A (green) and 0A (purple). Letters (a, b, or c) in the superscript represent the observed significant differences (Tukey’s test; P < 0.05).

### Putative electron donors for fixation of CO_2_ via the CBB cycle

Sulfite and hydrogen could be the putative electron donors of strain B3 because of the following evidence. Firstly, the strain harboured and transcribed the key genes *sox*YZ and those for [NiFe]-H_2_ases responsible for sulfite [68, 69] and H_2_ [70] oxidation to provide electrons, respectively (Table S3). Secondly, the cell biomass (dry weight) was remarkably improved by the addition of Na_2_SO_3_ or the increase in H_2_ as headspace gas, with the maximum increases of 2.82- and 1.12-fold in the presence of Na_2_SO_3_ and H_2_ in this experiment, respectively.

### Photosynthesis-related genes in strain B3 may be vertically evolved from its facultative relatives

G+C contents, codon usage, amino acid usage, and gene position were usually used to understand the gene’s origin, i.e., acquisition by inheritance or horizontal gene transfer (HGT) [71]. As described above, strain B3 harbours photosynthesis-related genes, including genes related to light utilization (clustered into PGCs as in other AAnPB) and CO_2_ fixation (in the CBB and CAM cycles). These genes had similar GC contents to that of the whole genome (Table S3). The similar GC content implied that the genes may be inherited in the genome and not inserted via HGT [72]. Moreover, BLAST analyses revealed that most of the photosynthesis-related genes of strain B3 were affiliated with genes from the Rhodospirillaceae family (Table S3). Therefore, it is most likely that the genes for light harnessing and the CO_2_ fixation pathways in strain B3 are vertically inherited and evolved rather than acquired through HGT from its facultative relatives.

The phylogenetic position of strain B3 further supported the above hypothesis, as the photosynthetic gene clusters (*puf*MLC-*bch*XYZ), *rbc*L and *rbcS* coding for RubisCO, and the 16S rRNA gene were most closely related to the corresponding genes in the Rhodospirillaceae family (Figure 6), which is distinct from the incongruences within the phylogenetic topologies between PGCs and 16S rRNA genes that represents a common incident of HGT of PGC in AAnPB [73]. The taxonomic position of strain B3 among purple phototrophic bacteria implied that the strain B3 probably evolved from an anaerobic ancestor of Rhodospirillaceae. Anaerobic members of Rhodospirillaceae had earlier differentiation times than close neighbours in Time Tree [74, 75]. During the adaptation to from anoxic to oxic conditions on Earth, anaerobic anoxygenic phototrophic bacteria evolved into AAnPB by modifying their photosynthetic apparatus biosynthesis strategy in response to oxygen levels, such as by the loss or acquisition of certain abilities [41]. However, as the cost of phototrophy is high compared to that in other presently known AAnPB, light energy is auxiliary to heterotrophy based on OC consumption [9]. Carbon fixation via the CCB cycle requires more energy [66], and no other pure-cultured AAnPB is known to use this pathway today. Surprisingly, strain B3 inherited not only a light energy utilization apparatus but also CO_2_ fixation by the CBB cycle from its anaerobic phototrophic ancestor, which suggests that the strain represents a novel AAnPB lineage.

**Figure 6.**
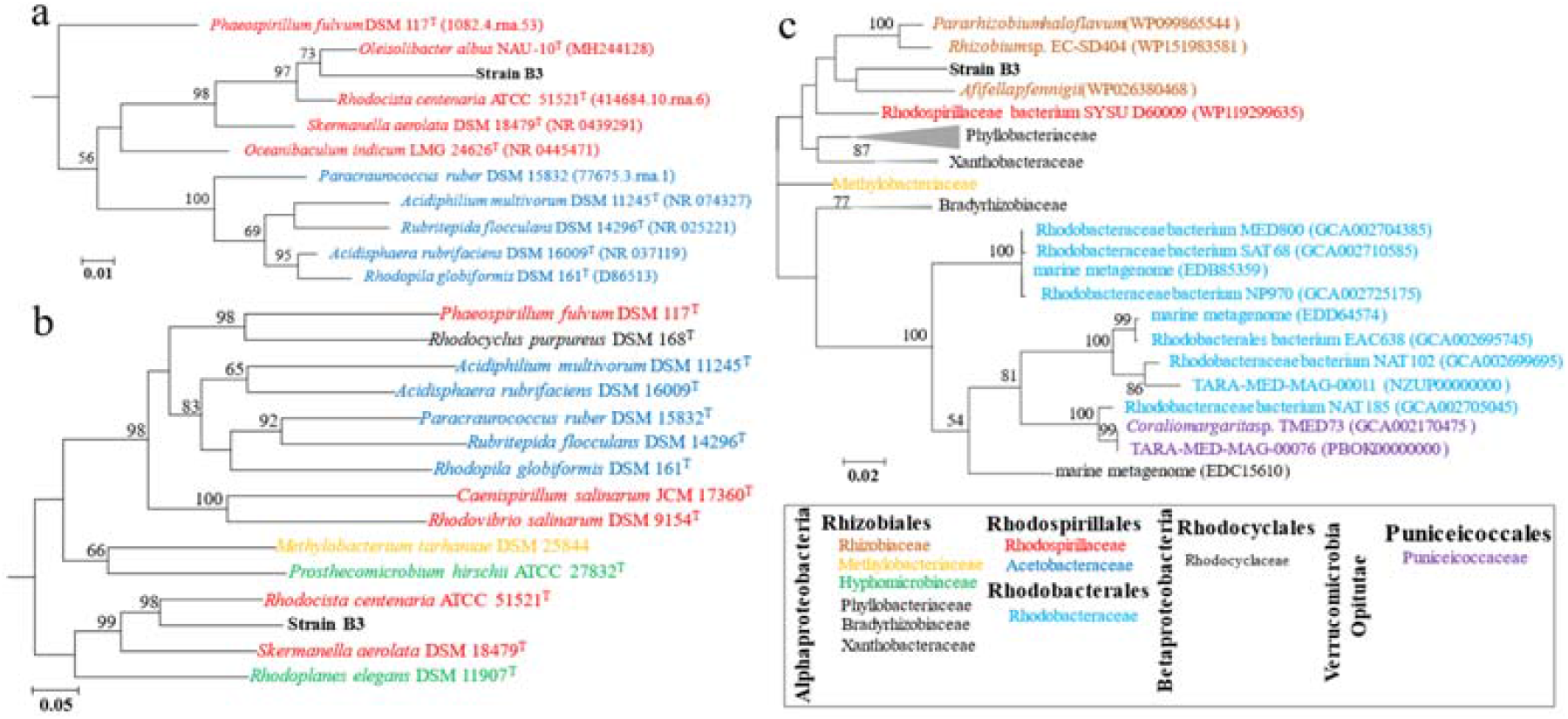
Phylogenetic tree based on 16S rRNA gene (a) sequences, *puf*MLC-bchXYZ gene clusters (b) and *rbc*L genes (c). A phylogenetic tree was constructed using both ML methods. Numbers at nodes are bootstrap values (ML, X, no bootstrap value in ML tree where nodes differ in both dendrograms; value<50% not shown). The bar represents 0.01, 0.02 and 0.05 substitutions per alignment position.

## Conclusion

Strain B3 represents a novel aerobic anoxygenic phototrophic lineage characterized by photoautotrophy using the Calvin-Benson-Bassham (CBB) cycle, in contrast to the photoheterotrophy of other currently known pure-cultured AAnPB. This kind of AAnPB may be ecologically significant in the global carbon cycle.

## Acknowledgements

This research was supported by grants from the National Natural Science Foundation of China (No. 31560030 & 31760025 & 32001220).

## Author contributions

KT carried out the experiments and prepared the draft of the manuscript. KT, YL, HL, BY, KJ and YZ analysed the data. FF and BY designed the research and wrote the manuscript with input from KT.

## Compliance with ethical standards

The authors declare that they have no conflicts of interest.

