## Supplemental Figure S1 for "An Aerobic Anoxygenic Phototrophic Bacterium Fixes CO_2_ via the Calvin-Benson-Bassham Cycle"

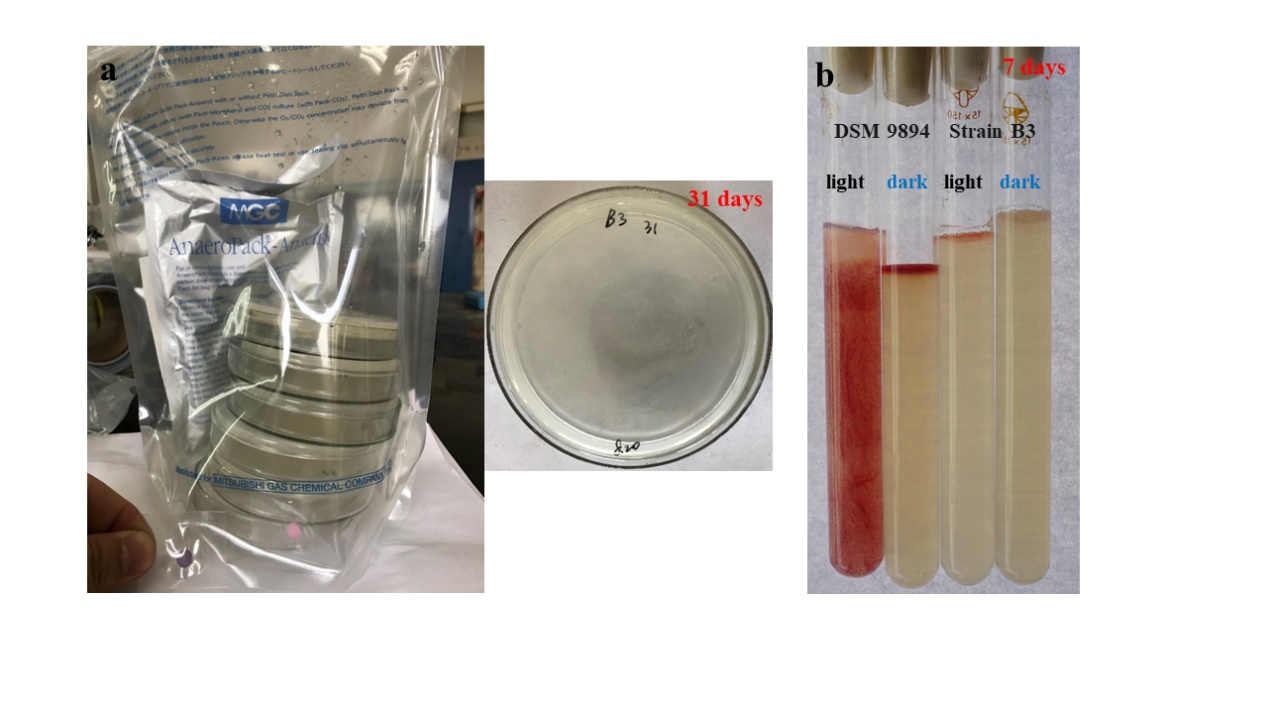

**Figure S1**. For the MGC AnaeroPack-Anaero (a), the oxygen indicator was red in anaerobic conditions. Strain B3 was cultured on 1/2-strength R2A agar medium in anaerobic packs for 31 days. Oxygen requirements were tested in a long tube (b) with Medium A.
