## Supplemental Figure S2 for "An Aerobic Anoxygenic Phototrophic Bacterium Fixes CO_2_ via the Calvin-Benson-Bassham Cycle"

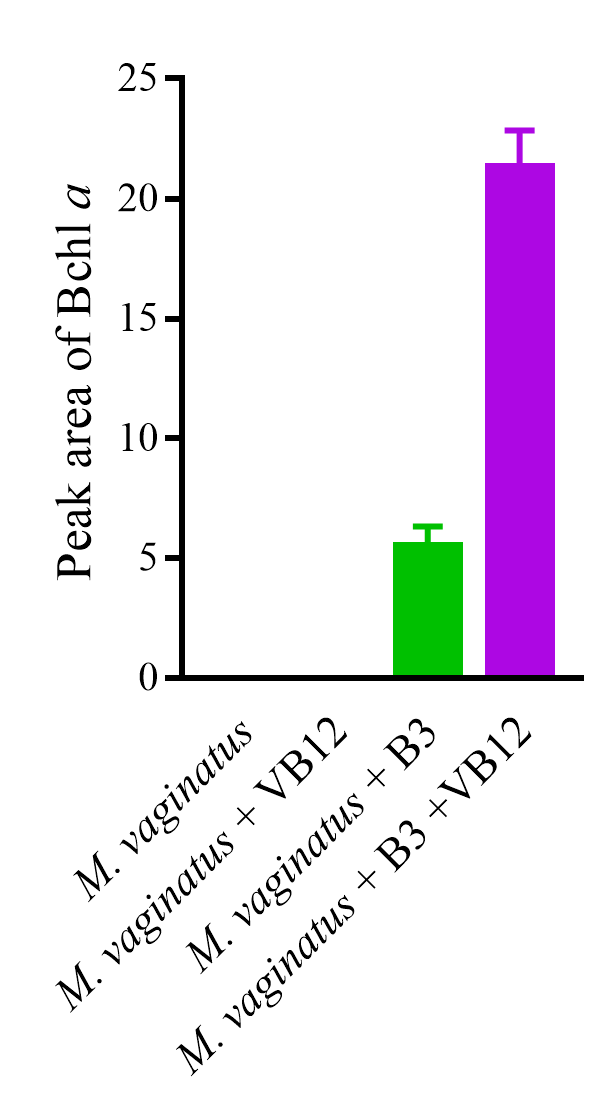

**Figure S2.** Bchl *a* was detected based on a coculture of strain B3 with *Microcoleus vaginatus* in liquid BG11. Cells of B3 and *M. vaginatus* were grown in 1/10 R2A and BG11 medium for five days, respectively. Then, the samples were centrifuged, washed three times and resuspended in NaCl solution (0.08%). One hundred microlitres of cell suspension (approximately 1.00 at OD600 nm) was inoculated into 50 mL of BG11 liquid media with 0.4 µg/50 mL of VB12 and cultured in a shaker at 160 r/min under natural light for 5 days. The Bchl *a* peak absorption at 710 nm in HPLC and the peak area were autocalculated by the manufacturer’s software (Agilent Technologies Inc., Waldbronn, Germany). Bchl *a* was not detected in the *M. vaginatus* and *M. vaginatus +* VB12 groups.
