## Supplemental Table S1 for "An Aerobic Anoxygenic Phototrophic Bacterium Fixes CO_2_ via the Calvin-Benson-Bassham Cycle"

**Table S1**. Compositions of media used in this study. For all media, the pH was adjusted to 7.2 before sterilization, and all media were sterilized at 121 °C for 20 min.

| **Components** | **1/2 strength** | **1/10 strength** | **5A** | **2.5A** | **A** | **0A** |
| --- | --- | --- | --- | --- | --- | --- |
| Yeast extract (g) | 0.25 | 0.05 | 0.05 | 0.05 | 0.05 | 0.05 |
| Proteose peptone (g) | 0.25 | 0.05 | 0.05 | 0.05 | 0.05 | 0.05 |
| Glucose (g) | 0.25 | 0.05 | 0.05 | 0.025 | 0.01 | 0 |
| Soluble starch (g) | 0.25 | 0.05 | 0.05 | 0.05 | 0.05 | 0.05 |
| Na-pyruvate (g) | 0.15 | 0.03 | 0.03 | 0.015 | 0.006 | 0 |
| K_2_HPO_4_ (g) | 0.15 | 0.03 | 0.03 | 0.03 | 0.03 | 0.03 |
| MgSO_4_·7H_2_O (g) | 0.025 | 0.005 | 0.005 | 0.005 | 0.005 | 0.005 |
| Vitamin B12 solution (mL)  (10 mg in 100 mL H_2_O) | 0.20 | 0.04 | 0.04 | 0.04 | 0.04 | 0.04 |
| Trace element solution SL-6 (mL)  (refer to DSMZ medium 27) | 0.50 | 0.10 | 0.10 | 0.10 | 0.10 | 0.10 |
| Distilled water (L) | 1.0 | 1.0 | 1.0 | 1.0 | 1.0 | 1.0 |

**BG11 medium:** citric acid (6 mg/L), ferric ammonium citrate (6 mg/L), EDTA-Na_2_ (1 mg/L), NaNO_3_ (0.6 g/L), K_2_HPO_4_ (15.6 mg/L), MgSO_4_·7H_2_O (30 mg/L), CaCl_2_·2H_2_O (38 mg/L), Na_2_CO_3_ (20 mg/L), H_3_BO_3_ (2.86 mg/L), ZnSO_4_·7H_2_O (0.222 mg/L), MnCl_2_·4H_2_O (1.81 mg/L), Na_2_MoO_4_ ·2H_2_O (0.391 mg/L), CuSO_4_·5H_2_O (0.079 mg/L), Co(NO_3_)_2_·6H_2_O (0.049 mg/L), vitamin B12 (8 μg/L), NiCl_2_·H_2_O (2.68 mg/L), Na_2_SO_3_ (48 mg/L).

**basal carbon-free Medium A** contained^*^ (g/l): MgSO_4_·7H_2_O (3), Na_2_SO_4_ (1.8), KH_2_PO_4_ (0.3), NH_4_Cl (0.3), KCl (0.7), CaCl_2_·2H_2_O (0.05), Difco Bactopeptone (0.5), Casamino Acids (0.5), Na_2_S_2_O_3_ (0.25), Na_2_SO_3_ (0.25), TES 2 ml, VS 2 ml.

Note: 1/2 strength R2A was used in primary isolate, 1/10 strength R2A in subculture for genome sequencing and BChl *a* detection; 5A, 2.5A, A and 0A were applied to reveal the effects of organic-carbon levels on the growth, photosynthetic genes expression, transcriptomics and RubisCo enzyme activity. basal carbon-free Medium A and BG11 medium was used in aerobic photoautotrophic growth tests.

^*^Csotonyi JT, Stackebrandt E, Swiderski J, Schumann P and Yurkov, V. An alphaproteobacterium capable of both aerobic and anaerobic anoxygenic photosynthesis but incapable of photoautotrophy: *Charonomicrobium ambiphototrophicum*, gen. nov., sp. nov. *Photosynth res* 2011;107: 257-268.
