## Supplemental Table S2 for "An Aerobic Anoxygenic Phototrophic Bacterium Fixes CO_2_ via the Calvin-Benson-Bassham Cycle"

**Table S2.** PCR primers and thermal cycling conditions used for the amplification and quantification of different genes.

| **Gene designation** | **Representatives of various taxa** | **Primer sequences (5'-3')** | **Temperature profile** | **Standard curve** |
| --- | --- | --- | --- | --- |
| **16S rDNA^1, 2^** 27F 1492R | Bacteria | AGAGTTTGATCMTGGCTCAG GGTTACCTTGTTACGACTT | 95 °C, 5 min, 1 cycle  95 °C for 30 s, 55 °C for 45 s, 72 °C for 90 s, 35 cycles 72 °C for 10min |  |
| **16S rDNA^3^** 1369F 1541R | Bacteria | CGGTGAATACGTTCYCGG AAGGAGGTGATCCRGCCGCA | 95 °C, 1 min, 1 cycle  95 °C for 15 s, 58 °C for 20 s, 72 °C for 30 s, 40 cycles 95 °C for 15 s, 60 to 95 °C, 1 cycle | y=-0.3196x+13.578 R^2^=0.9996 |
| ***puf*M^4^** *puf*M 557F *puf*M_WAW | AAP bacteria | TACGGSAACCTGTWCTAC CCATSGTCCAGCGCCAGAA | 95 °C, 30s,1 cycle  95 °C for 10 s, 52 °C for 15 s, 72 °C for 30 s, 40 cycles 95 °C for 15 s, 60 to 95 °C, 1 cycle | y=-0.3137x+12.504  R^2^=0.9982 |
| ***rbc*L^5, 6^**  k2f v2R | Bacteria with RubisCO | ACCAYCAAGCCSAAGCTSGG GCCTTCSAGCTTGCCSACCRC | 95 °C, 3 min, 1 cycle  95 °C for 30 s, 58 °C for 30 s, 72 °C for 30 s, 40 cycles 95 °C for 30 s, 60 to 95 °C, 1 cycle | y=-0.2876x+12.503  R^2^=0.9963 |
