## Supplemental Table S4 for "An Aerobic Anoxygenic Phototrophic Bacterium Fixes CO_2_ via the Calvin-Benson-Bassham Cycle"

**Table S4.** Basic information on the strain B3 genome.

| Category | Name | Chromosome | Plasmid A | Plasmid B | Plasmid C | Plasmid D |
| --- | --- | --- | --- | --- | --- | --- |
| Basic information | No. of all scaffolds | 1 | 1 | 1 | 1 | 1 |
|  | Total length (bp) | 3664154 | 935201 | 712893 | 702078 | 190713 |
|  | G+C content (mol %) | 69.264 | 72.609 | 71.268 | 70.552 | 75.364 |
|  | N rate (%) | 0 | 0 | 0 | 0 | 0 |
| Gene information | Gene numbers | 3614 | 873 | 614 | 631 | 193 |
|  | Gene total length (bp) | 3160236 | 804327 | 632820 | 595968 | 146298 |
|  | Gene average length (bp) | 841 | 921 | 1030 | 944 | 758 |
|  | Gene density (No./kb) | 1.025 | 0.933 | 0.861 | 0.898 | 1.011 |
|  | GC content in gene region (%) | 69.7 | 73 | 71.8 | 71.3 | 75.8 |
|  | Gene/Geonme (%) | 86.2 | 86 | 88.8 | 84.9 | 76.7 |
| RNA prediction | No. of tRNA (types) | 48 (19) | 4 (4) | 1 (1) | 1 (1) | 0 |
|  | No. of 5S rRNA | 2 | 1 | 0 | 2 | 0 |
|  | No. of 16S rRNA | 2 | 1 | 1 | 2 | 0 |
|  | No. of 23S rRNA | 2 | 1 | 1 | 2 | 0 |
