## Supplemental Table S5 for "An Aerobic Anoxygenic Phototrophic Bacterium Fixes CO_2_ via the Calvin-Benson-Bassham Cycle"

**Table S5.** Pearson correlation between added carbon contents, B3 cell parameters and different gene copy numbers.

| Variable | Carbon  add content | Cell | | | Copy numbers (log10) | | |
| --- | --- | --- | --- | --- | --- | --- | --- |
|  |  | size | density | biomass | 16S rRNA | *puf*M | *cbb*L |
| Carbon add content |  |  |  |  |  |  |  |
| Cell size | -0.17 |  |  |  |  |  |  |
| Cell density | 0.93 | -0.45 |  |  |  |  |  |
| Cell biomass | 0.88 | -0.59 | 0.99* |  |  |  |  |
| 16S rRNA | 0.87 | -0.53 | 0.99* | 0.99* |  |  |  |
| *puf*M | 0.95 | -0.37 | 0.99** | 0.97* | 0.98* |  |  |
| *cbb*L | 0.63 | -0.87 | 0.8 | 0.89 | 0.83 | 0.75 |  |

Note: * and **Correlation is significant at the 0.05 and 0.01 level (two-tailed), respectively.
